# Phylogenetic Network Dissimilarity Measures That Take Branch Lengths Into Account

**DOI:** 10.1101/2022.02.22.481483

**Authors:** Berk A. Yakici, Huw A. Ogilvie, Luay Nakhleh

## Abstract

Dissimilarity measures for phylogenetic trees have long been used for analyzing inferred trees and understanding the performance of phylogenetic methods. Given their importance, a wide array of such measures have been developed, some of which are based on the tree topologies alone, and others that also take branch lengths into account. Similarly, a number of dissimilarity measures of phylogenetic networks have been developed in the last two decades. However, to the best of our knowledge, all these measures are based solely on the topologies of phylogenetic networks and ignore branch lengths. In this paper, we propose two phylogenetic network dissimilarity measures that take both topology and branch lengths into account. We demonstrate the behavior of these two measures on pairs of related networks. Furthermore, we show how these measures can be used to cluster a set of phylogenetic networks obtained by an inference method, illustrating this application on the posterior sample of phylogenetic networks. Both measures are implemented in the publicly available software package PhyloNet.

## 1 Introduction

Phylogenetic trees and networks are widely used to model the evolution of genes, species and languages. In the case of genomes and species, a deluge of data is available in the form of whole genomes being assembled and made available for phylogenetic inference [15, 20, 25]. However, the space of phylogenetic trees is notoriously complex due to a mix of discrete and continuous parameters. Therefore, this complexity must be confronted, as probability and likelihood distributions over phylogenetic trees often lack closed form solutions, requiring exploration of this space using algorithms such as hill climbing or Markov-chain Monte Carlo. Worse still, the discrete and continuous parameters are highly dependant, making this exploration fiendishly difficult. Phylogenetic network distributions are more complex again, among other reasons because dimensionality of the problem is no longer fixed.

Nonetheless, phylogenetic trees and networks remain successful models due to their strong and direct relationship to actual evolution. Evolving units such as species split and diverge over time, and patterns of splits are encoded by the nodes of phylogenetic trees. Each node represents a species (or other kind of evolving unit), and has one parent and a certain number of children (typically zero or two) to connect with its immediate ancestor and descendants respectively. Units which we have data for are typically represented as external nodes, often called taxa for trees of species or populations. The degree of divergence can be encoded as either the branch lengths or, for ultrametric phylogenies, as node ages. The pattern of splits and reticulations without continuous parameters is known as the tree topology and is an unordered discrete variable that grows hyperexponentially with the number of taxa [10].

Because phylogenetic trees do not account for reticulate evolution, such as introgression, hybridization, or horizontal gene transfer between species, or loanwords borrowed between languages [13], phylogenetic networks were developed in order to model splitting and reticulation. These networks may contain reticulation nodes, with two parents and one child, in addition to the nodes used to encode splits. The probability of inheriting evolutionary subunits, such as genes or words, may be encoded as additional continuous parameters associated with the reticulation nodes. By permitting two immediate ancestors with inheritance probabilities, phylogenetic networks can effectively encode the aforementioned examples of reticulate evolution [9, 21]. The addition of reticulation nodes makes the number of possible phylogenetic network topologies far greater than the number of trees. Deriving these numbers are non-trivial, and so far have been restricted to specific classes of networks. For level-2 networks, there are 1,143 network topologies for three external nodes, compared with only three tree topologies [5].

The complexity of phylogenetic tree and network space has some immediate implications. For example, to quantify the dissimilarity between sets of continuous parameter values we can choose among the *L^p^*-norms (e.g. the Euclidean norm), but this is not directly applicable to phylogenies due to their complex mixture of discrete and continuous parameters that cannot naturally be embedded in *L^p^* spaces. However, we often wish to measure this dissimilarity to compare different inference methods with each other, to compare them with ground truths, or to study how different sets of evolutionary units co-evolve [2]. Furthermore, novel efficient proposal kernels such as Zig-Zag cannot be applied to phylogenetic trees without thorough preliminary theoretical work [17]. The Zig-Zag traverses tree space, randomly reversing direction along a given dimension at intervals. It is not obvious how a particle can sample multiple tree topologies by proceeding along or reversing the direction of travel, although some preliminary work has been done in this area [8].

These implications have motivated the development of myriad measures and metrics of dissimilarity and distances to compare phylogenetic trees, which have enabled the comparison of different phylogenies and improved our understanding of the algorithms used in phylogenetic inference [34]. We can classify many of these measures and metrics into three broad categories: clade-based, move-based, and geometric.

Clade-based measures are based on the presence or absence of clades (or, for unrooted phylogenies, splits). These may be limited to the difference in topology as in the Robinson-Foulds (RF) distance [26], or incorporate branch lengths as in weighted RF and branch score (BS) distances [18]. Move-based measures are based on the number of random-walk moves, such as Nearest-Neighbor Interchange (NNI), needed to modify one topology to be identical with another topology [1]. Geometric measures rely on the embedding of phylogenies in geometric spaces, such as the Billera-Holmes-Voghtmann (BHV), *τ* and *t* spaces [3, 11]. The distance between phylogenies is then the shortest path between the corresponding points.

The additional complexity of phylogenetic networks means that available measures and metrics are far less developed. However, the need to accurately model evolution with reticulation demands much greater development and motivates our present study. Presently, beyond simple identity, there are several measures of topological dissimilarity with proofs of constituting metrics on subspaces of phylogenetic networks, e.g., [6, 7, 22].

None of these existing measures considers branch lengths or node ages, despite the importance of these distances in evolutionary biology. To address this absence, in this paper we propose rooted network branch score (rNBS) and average path distance (APD), two novel measures to compute dissimilarity between two rooted phylogenetic networks *Ψ*_1_ and *Ψ*_2_ that are sensitive to branch lengths in addition to the topology. When comparing pairs of simulated networks that undergo reticulation elimination and branch-length scaling, we observe an increase in dissimilarity value from both measures with respect to the amount of distortion applied to one of the pairs. Additionally, when we use rNBS, APD, and the topological distance of [22] to cluster networks obtained from a Bayesian Markov chain Monte Carlo (MCMC) sample based on their dissimilarity, we find that both rNBS and APD can highlight structure within searches of tree space that may not be obvious from other parameters and statistics. Thus, we believe both rNBS and APD are suitable in evaluating, prototyping and refining network inference. Both measures are implemented in PhyloNet [31, 33] and are publicly available to download and use. The source code is available at https://github.com/NakhlehLab/PhyloNet/.

## 2 Methods

Our focus in this paper is binary evolutionary (or, explicit) phylogenetic networks. Furthermore, we assume all networks are leaf-labeled by the same set of taxa.

### Definition 1.

*The topology of a phylogenetic network Ψ is a rooted directed acyclic graph (V, E) such that V contains a unique node with in-degree* 0 *and out-degree* 2 *(the root) and every other node has either in-degree* 1 *and out-degree* 2 *(an internal tree node), in-degree* 1 *and out-degree* 0 *(an external tree node, or leaf), or in-degree* 2 *and out-degree* 1 *(a reticulation node). Edges incident into reticulation nodes are referred to as reticulation edges. The leaves are bijectively labeled by a set* 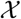 *of taxa, with* 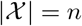. *The phylogenetic network Ψ has branch lengths* 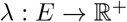, *where λ_b_ denotes the branch length of branch b in Ψ*.

In this section, we propose two different methods of measuring the dissimilarity of a pair of phylogenetic networks *Ψ*_1_ and *Ψ*_2_ while taking their branch lengths into account.

### 2.1 Rooted Network Branch Score (rNBS)

In this subsection, we describe a dissimilarity measure based on viewing a network in terms of the trees it displays, similar to the tree-based measure for topological comparison of phylogenetic networks implemented in PhyloNet [31].

#### Definition 2.

*Let Ψ be a phylogenetic network leaf-labeled by set* 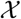 *of taxa. A tree T is displayed by Ψ if it can be obtained from Ψ by removing for each reticulation node exactly one of the edges incident into it followed by repeatedly applying forced contractions until no nodes of in- and out-degree* 1 *or in-degree* 0 *and out-degree* 1 *remain. A forced contraction of a node u of in-degree* 1 *and out-degree* 1 *consists of (i) adding an edge from u’s parent to u’s child, and (ii) deleting node u and the two edges that connect it to its parent and child. A forced contraction of a node u of in-degree* 0 *and out-degree* 1 *consists of removing the node u and its incident edge. The resulting tree has a unique root, whose in-degree is* 0 *and out-degree is* 2, *leaf nodes, whose in-degrees are* 1 *and out-degrees are* 0, *and other internal nodes, whose in-degrees are* 1 *and out-degrees are* 2. *We denote by* 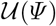 *the set of all trees displayed by Ψ, where each tree in* 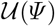 *is leaf-labeled by set* 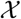 *of taxa; that is, no tree whose leaves are not bijectively labeled by* 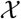 *is included in the set* 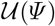.

It is important to note here that since branch lengths are taken into account, the set 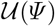 can have trees with identical topologies but different branch lengths. This is illustrated in Fig. 1.

**Fig. 1:**
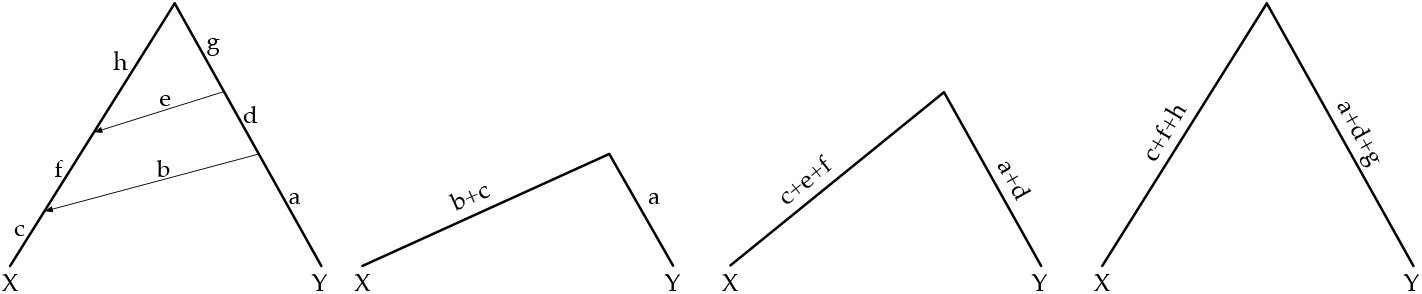
A phylogenetic network (left) and its displayed trees. Since branch lengths are taken into account, the network displays three different trees. In the case of topology alone, the network displays a single tree.

We build a weighted complete bipartite graph 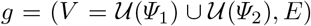 where the weight of edge (*t*_1_, *t*_2_) ∈ *E* equals the rooted branch score of *t*_1_ and *t*_2_ [12], which is the rooted equivalent of the branch score of [18]. The rNBS of *Ψ*_1_ and *Ψ*_2_ is then computed as the minimum-weight edge cover of *g* normalized by the number of edges in the edge cover. An edge cover of *g* is a subset *E*′ ⊆ *E* of its edges so that every node in *V* is the endpoint of at least one edge in *E*′. The weight of an edge cover *E*′ is the sum of the weights of the edges in *E*′. A minimum-weight edge cover is an edge cover of *g* whose weight is smallest among all possible edge covers of *g*.

In our implementation, we use the Hungarian method to compute edge cover, which runs in 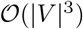. However, in its current implementation, rNBS is not scalable with respect to the number of reticulations, since the size of V grows exponentially in the number of reticulation nodes. Exploring whether the rNBS value between two networks can be computed more efficiently without explicitly building the bipartite graph is a direction for future research. Fig. 2 illustrates the rNBS of two phylogenetic networks.

**Fig. 2:**
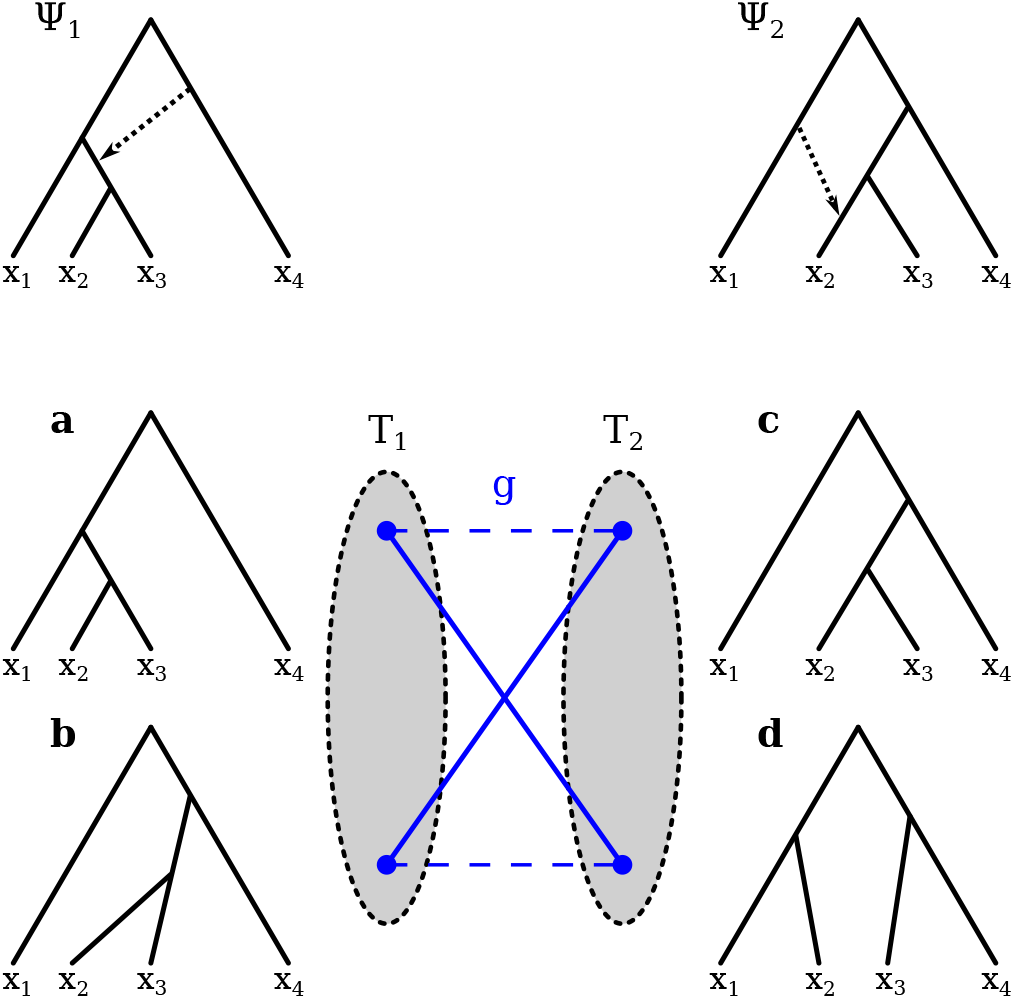
Illustrating the rooted network branch score (rNBS) of two phylogenetic networks *Ψ*_1_ and *Ψ*_2_. 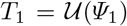 and consists of the two trees (**a**-**b**) with distinct branch lengths and 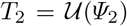 and consists of the two trees (**c**-**d**) with distinct branch lengths. The complete bipartite graph *g* = (*T*_1_ ∪ *T*_2_, *E*) is shown. Assuming the minimum-weight edge cover consists of the two edges depicted by the blue solid lines, then rNBS(*Ψ*_1_, *Ψ*_2_) = *w*(**a**, **d**) · 0.5 + *w*(**b**, **c**) · 0.5.

Additionally, it is important to note that the rNBS is not a metric, in particular failing to satisfy the condition that rNBS(*Ψ*_1_, *Ψ*_2_) = 0 if and only if *Ψ*_1_ and *Ψ*_2_ are isomorphic (while respecting the leaf labeling). Fig. 3 shows two networks that display the same set of trees even when branch lengths are included. One setting of the network branch lengths that would lead to this scenario is given by the following Rich Newick strings of the two networks:

Psi1 = (((((X2:7.0)#H2:4.0)#H1 : 6.0, X3:8.0) : 4.0, (#H1 : 4.0, X4 : 5.0) : 2.0) : 3.0, (#H2:2.0, X1 : 2.0) : 1.0) Root;
Psi2 = ((((X2:4.0)#H1 : 13.0, X3 : 8.0) : 4.0, ((#H1:3.0)#H2 : 8.0, X4:5.0) : 2.0) : 3.0, (#H2 : 2.0, X1 : 2.0) : 1.0) Root;

**Fig. 3:**
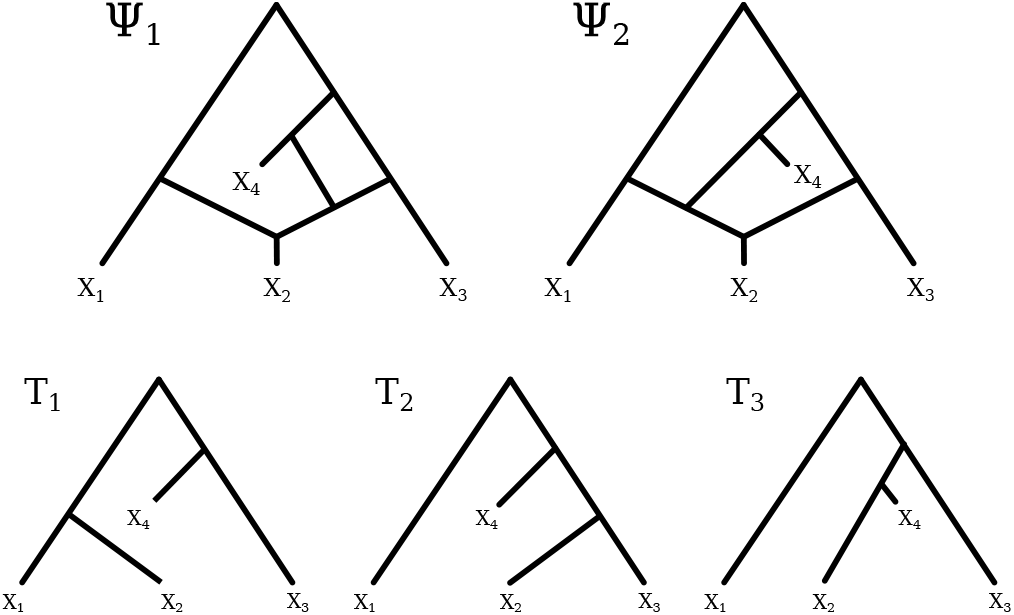
Two networks *Ψ*_1_ and *Ψ*_2_ that display the same set of trees {*T*_1_, *T*_2_, *T*_2_} (adapted from [23]).

In this case, rNBS(*Ψ*_1_, *Ψ*_2_) = 0 even though the two networks are different.

### 2.2 Average Path Distance (APD)

Here we present a dissimilarity measure that is based directly on the networks, rather than the trees they display. We view phylogenetic networks *Ψ*_1_ and *Ψ*_2_ as two *n* × *n* matrices *M*_1_ and *M*_2_, respectively, where entries [*i, j*] in *M*_1_ and *M*_2_ correspond to the average path distance between the two leaves labeled i and *j* in *Ψ*_1_ and *Ψ*_2_, respectively. Thus, it can be viewed as an extension of path distance by [28] to networks.

In a phylogenetic tree, the path distance between two leaves is the sum of weights of edges on the unique (simple) path between those two leaves. In a phylogenetic network, there could be more than one path between two leaves. Let 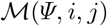 be the set of all most recent common ancestors (MRCAs) of *i* and *j* in network *Ψ*. Here, an MRCA is a node from which there is a path to *i* and a path to *j* and these two paths do not share any edge. The average path distance (APD) between two leaves *i* and *j* is the average of all such paths between the two leaves. For example, in the network shown in Fig. 4, there are three MRCAs of *X*_2_ and *X*_3_.

**Fig. 4:**
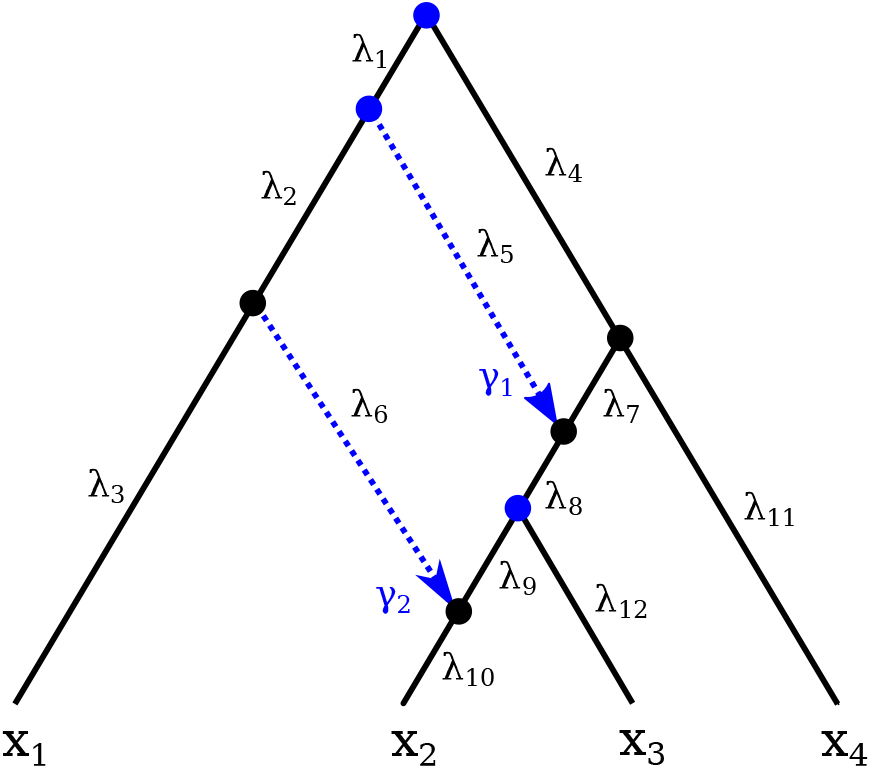
Illustrating the average path distance (APD) on a phylogenetic network. The blue solid circles correspond to the three MRCAs of *X*_2_ and *X*_3_. The distances between *X*_2_ and *X*_3_ that go through these three MRCAs are: (λ_10_ + λ_6_ + λ_2_ + λ_1_) + (λ_12_ + λ_8_ + λ_7_ + λ_4_) (through the root node as the MRCA), (λ_10_ + λ_6_ + λ_2_) + (λ_12_ + λ_8_ + λ_5_) (through the MRCA that is a child of the root), and (λ_10_ + λ_9_) + λ_12_ (through the MRCA that is the parent node of *X*_3_). The average path distance (APD) of *X*_2_ and *X*_3_ is the average of these three distances.

To compute matrices *M*_1_ and *M*_2_ we utilize a BFS-like approach to traverse networks *Ψ*_1_ and *Ψ*_2_, starting from their leaves and finishing at the root, visiting each internal node if and only if we have explored all of its children first. If we are currently at an MRCA of leave pairs *i* and *j*, we add their path distance to matrix entries [*i, j*] and [*j, i*]. After traversing the graph, we divide each entry in the matrices by the number of MRCAs per pair of leaves to obtain the average path distance. After building matrices *M*_1_ and *M*_2_ for networks *Ψ*_1_ and *Ψ*_2_, respectively, APD can be computed by taking the Frobenius norm of the matrix difference:

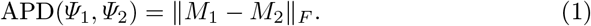

Overall, the graph traversal runs in 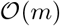, where *m* is the number of edges in the network. Building and summarizing each matrix takes 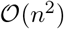. The computation in Eq. 1 also takes 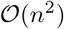 operations. Thus, APD is computable in polynomial time with respect to the number of nodes in the phylogenetic network.

However, it is important to note that the APD of the two different networks of Fig. 3 is also 0 under the branch length settings above. Therefore, APD is not a metric either.

#### Normalized average path distance (NormAPD)

We can obtain a normalized APD measure (NormAPD) as follows, which assumes *Ψ*_1_ is a reference network and is useful in settings where an inferred network is compared to a reference one.

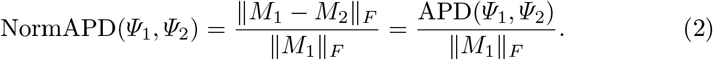

## 3 Results and Discussion

### 3.1 Dissimilarity Under Various Network Perturbations

To study the behavior of our dissimilarity measures on pairs of related networks, we simulated rooted phylogenetic networks that are leaf-labeled by the same set of 8 taxa using the SpeciesNetwork [35] add-on in BEAST 2.5 [4]. For the simulation, we set the origin to 0.1, birth rate to 30, and hybridization rate to 5. We then filtered out networks that contain more than 7 reticulations to limit the number of trees displayed by the networks. Our final data set contained 500 rooted phylogenetic networks. For each network *Ψ*_0_ in the set of simulated networks, we generated perturbed versions of *Ψ*_0_ and calculated the dissimilarity between the perturbed networks and *Ψ*_0_.

#### Uniform scaling

Here we obtain *Ψ_i_* by scaling each branch of *Ψ*_0_ by a factor of 1.5 for 10 iterations. At the end of iteration *i*, we have network *Ψ_i_* whose branch lengths are (1.5)^*i*^λ_0_, where λ_0_ is the branch length of the particular branch on the original network *Ψ*_0_. We computed rNBS(*Ψ*_0_, *Ψ_i_*) and APD(*Ψ*_0_, *Ψ_i_*), for *i* = 1, 2, …, 10, and plotted the results as a function of the iteration number. The results for rNBS and APD are shown in Figs. 5a and 5b, respectively. Both figures show an exponential relationship between the number of iterations and the dissimilarity between the original and perturbed networks measured by rNBS and APD. Furthermore, we observe that neither of the measures have a consistent rate of increase in the dissimilarity values across pairs of networks. In fact, without normalization, both measures report higher values of dissimilarity when the network contains more edges.

**Fig. 5:**
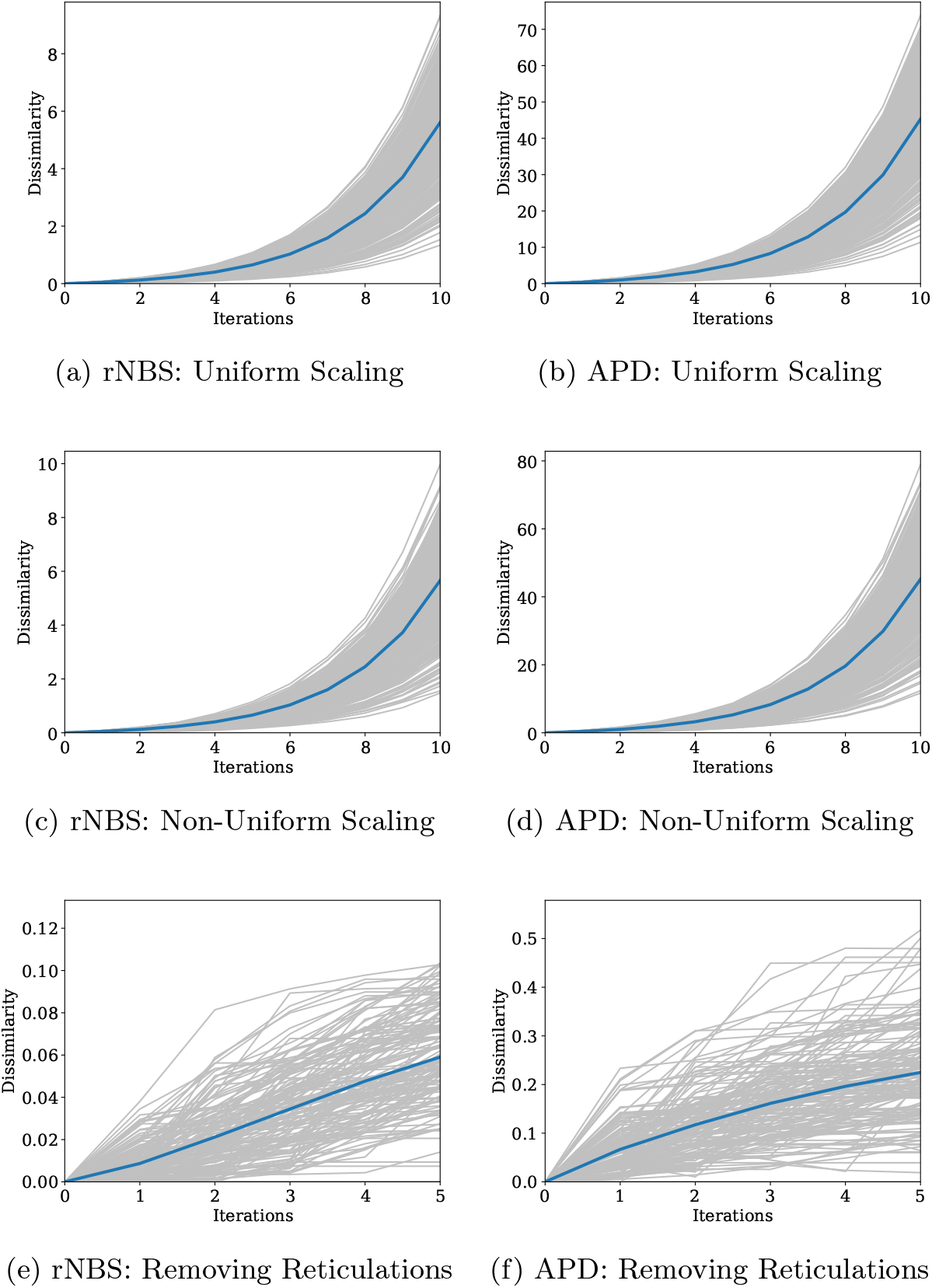
Dissimilarity measures on pairs of networks using three different perturbations. (a-b) Perturbed networks are obtained by uniform scaling of all branches and comparing perturbed networks to a reference network. (c-d) Perturbed networks are obtained by non-uniform scaling of all branches and comparing perturbed networks to a reference network. (e-f) Perturbed networks are obtained by removing reticulation edges and comparing perturbed networks to a reference network. Each gray line shows the relationship between number of iterations and the amount of dissimilarity between the original and perturbed networks. Blue lines show the average dissimilarity value at each iteration.

#### Non-uniform scaling

Here, we obtain *Ψ_i_*, for *i* = 1, 2, …, 10 from *Ψ*_0_ in a similar fashion to what we did in the case of uniform scaling, except that in each of the 10 iterations, we scaled each branch by a value drawn from 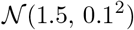. The results for rNBS and APD are shown in Figs. 5c and 5d, respectively. Even though scale factors are sampled from a distribution, the rates of increase of dissimilarity values computed by both rNBs and APD are consistent with the results from uniform scaling.

#### Reticulation elimination

Here, we further filter out networks that have fewer than 5 reticulation nodes, thus limiting our dataset to networks with at least 5 and at most 7 reticulations, which accounts for 157 networks. We produce *Ψ_i_* at each iteration as follows: *Ψ*_1_ is obtained by removing a random reticulation edge from *Ψ*_0_, and then *Ψ_i_* is obtained by removing a random reticulation edge from *Ψ*_*i*–1_, for *i* = 2, …, 5. The results for rNBS and APD are shown in Figs. 5e and 5f, respectively.

As the figure shows, the dissimilarity values increase as more reticulations are removed, but that increase slows down for the APD values with the number of removed reticulations, which is apparent in Fig. 5f. For the rNBS values (Fig. 5e), observe that the change in values is slow as the average value goes from 0 to around 0.06 when 5 reticulations are removed. The reason for this is that while reticulations are removed, the rest of the topologies and branch length are unperturbed. Therefore, it is natural that the removal of reticulations would have much less of an effect than, say, having all branches differ in lengths between the reference and perturbed networks.

In summary, perturbation experiments show that the measures we introduce here are sensitive to both changes in branch lengths and reticulation edges and behave as expected from dissimilarity measures.

### 3.2 Analyzing Posterior Samples Using the Dissimilarity Measures

Phylogenetic analyses oftentimes produce thousands of candidates. To summarize the tree candidates, a consensus tree is often computed, e.g., [16, 30]. As an alternative to a single consensus tree, clustering was offered as early as 1991 by W. Maddison [19]. Subsequently, it was shown that clustering indeed provides a more powerful and informative summary than single consensus trees [29]. Here, we explore the use of our dissimilarity measures for clustering networks in the posterior sample of a Bayesian inference method. We also compare them to clustering based on the topological dissimilarity measure of [22], referred to hereafter as TD.

MCMC_SEQ [32] is a method in PhyloNet [31, 33] for Bayesian inference of phylogenetic network topology, divergence times, and inheritance probabilities, along with various other parameters from sequence alignment data. The method samples from the posterior distribution of these parameters. We analyzed the posterior samples obtained by MCMC_SEQ on one simulated data set and one empirical data set.

#### Analysis of a simulated data set

We analyze the posterior sample obtained on a simulated data set of 5 species and a single individual per species. To generate the data set, we first simulated the homogeneous gene trees with MS [14], obtaining 100 gene trees with 500 sites per gene tree. Finally, we use Seq-Gen [24] to generate the simulated sequences. For this, we set the base frequencies {*A*, *C, G, T*} = {0.2112, 0.2888, 0.2896, 0.2104} and theta to 0.018. We also varied the substitution rates to follow a flat Dirichlet distribution. The sequences are then inputted to MCMC_SEQ, setting the Markov chain Monte Carlo (MCMC) chain length to 50, 000, 000, the burn-in period to 10, 000, 000, and the sample frequency to 5000.

The networks in the posterior sample vary in their number of reticulation nodes between 0 (i.e., trees) and 4. However, if we only consider the samples after burn-in period is completed (that is, ignoring the first 2,000 samples), the number of reticulations alternate between 3 and 4. Since clustering would require computing all pairwise distances between the networks in the sample, we further reduced the sample by keeping only every 5th sample, i.e., the 2000th, 2005th, …, and 9995th samples (the first 2,000 samples were discarded as part of the burn-in period). We then computed pairwise dissimilarity matrices using rNBS, APD, and TD of [22]. Afterwards, we clustered the posterior networks in two different ways:

1. Clustering I: We partitioned the samples based on the number of reticulation nodes in the networks, thus resulting in two clusters, one consisting of all networks with 3 reticulations and another consisting of all networks with 4 reticulations.
2. Clustering II: We applied agglomerative clustering to the pairwise distance matrices, and set the linkage criterion to “complete,” which makes the clustering algorithm use the maximum distances between all observations of the two sets when merging pairs of clusters. To determine the number of clusters, we looked at the dendrogram from each figure and manually curated the number to split. Note that cluster labels are arbitrarily assigned independently for each plot.

As the number of reticulations in networks is a major distinguishing factor when contrasting networks inferred on biological data, the rationale of the two ways of clustering is to understand (i) how the different iterations of MCMC_SEQ correlated in terms of the clustering, and (ii) whether clustering II is a refinement of clustering I.

For visualization, we plot the log-posterior density per iteration, number of reticulations per iteration (which corresponds to clustering I), the assigned cluster per iteration (which corresponds to clustering II), and the pairwise dissimilarity matrix computed by rNBS, APD, and TD of [22]. Here, we did not visualize the clustered heatmaps, but rather kept them ordered according to the sampling order since we are focused on understanding the correlation, in terms of dissimilarity, between adjacent samples. The results are shown in Fig. 6.

**Fig. 6:**
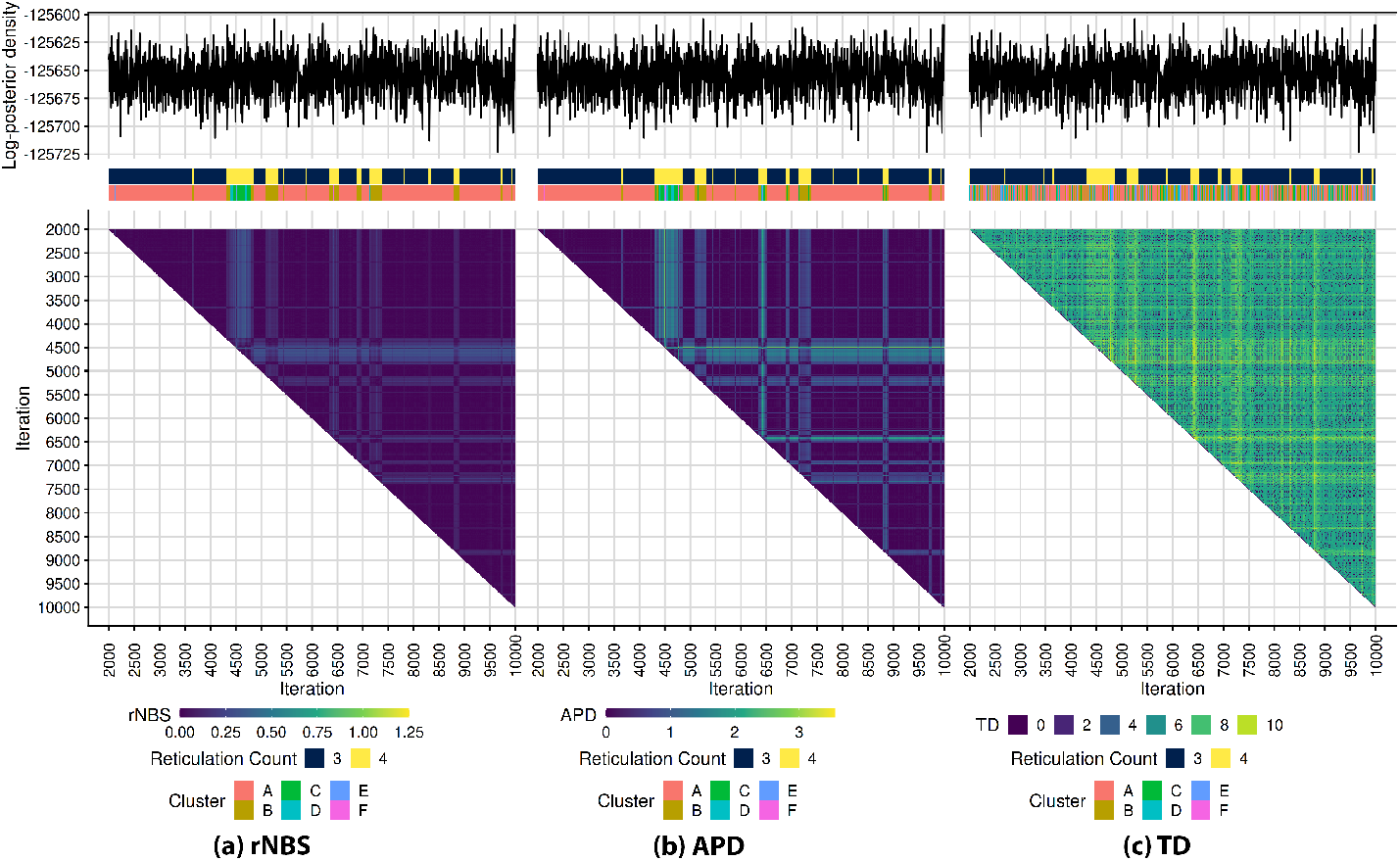
Clustering of samples (networks and their branch lengths) from the posterior distribution obtained by MCMC_SEQ on a simulated data set. Each panel shows the log-posterior density, the number of reticulations in each sampled network, the assigned clustering label obtained by agglomerative clustering, and the heatmap of the dissimilarity values arranged by the number of MCMC iterations. Panels (a)—(c) correspond to the three dissimilarity measures rNBS, APD, and TD, respectively.

From Fig. 6 we can observe that rNBS, APD, and TD all cluster the networks with respect to the number of reticulations, shown by the purple vs blue shades in Fig. 6 (a) for rNBS, purple vs green shades in Fig. 6 (b) for APD, and green vs yellow shades in Fig. 6 (c) for TD. While all heat-maps support the clustering by the number of reticulations, the clusters are best defined based on APD pairwise distances, followed by rNBS pairwise distances, and then least defined based on TD pairwise distances.

Additionally, agglomerative clustering obtained based on rNBS and APD pairwise distances shows distinct sub-clusters within the two clusters based on the number of reticulations alone. For example samples in two distinct clusters *B* and *C* for both rNBS and APD contain samples with 4 reticulations, suggesting cluster analyses can help discern patterns of similarity in terms of the posterior values and obtain a refined clustering of the posterior sample. The fact that these structures were only sampled in one segment of the posterior chain also suggests it is less converged than the log-posterior probability trace suggests, and that APD- and rNBS-based clustering are more powerful diagnostic tools for convergence.

#### Analysis of an empirical data set

For the empirical results, we analyzed the yeast data set of [27] with seven *Saccharomyces* species. We utilized the same methods as in the previous section, with the exception of setting the MCMC chain length to 35, 000, 000, obtaining 5, 000 samples after discarding the first 2,000. The results are shown in Fig. 7.

**Fig. 7:**
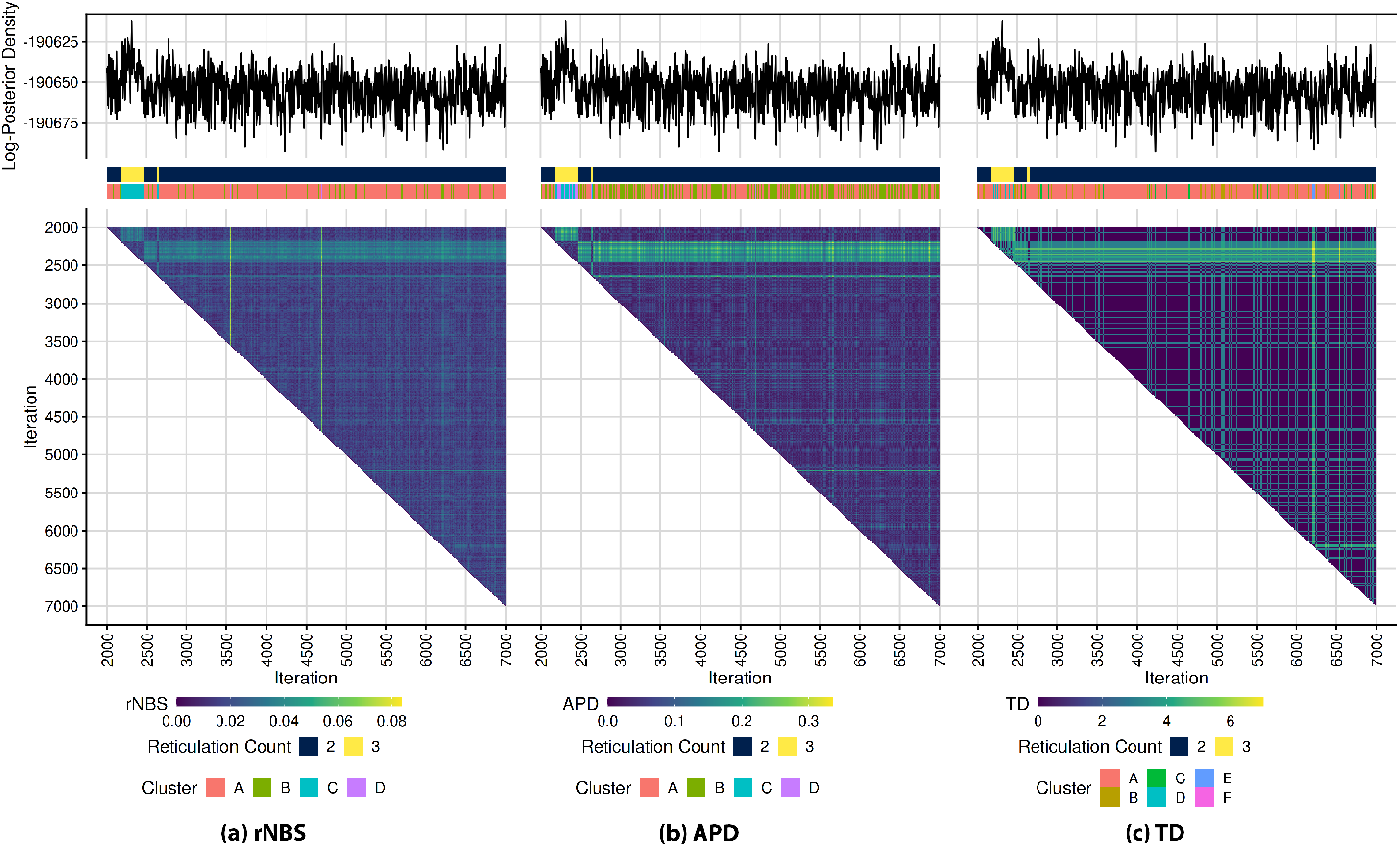
Clustering of samples (networks and their branch lengths) from the posterior distribution obtained by MCMC_SEQ on an empirical yeast data set. Each panel shows the log-posterior density, the number of reticulations in each sampled network, the assigned clustering label obtained by agglomerative clustering, and the heatmap of the dissimilarity values arranged by the number of MCMC iterations. Panels (a)—(c) correspond to the three dissimilarity measures rNBS, APD, and TD, respectively.

From Fig. 7, we see that both rNBS and APD highlight similar pairwise dissimilarity characteristics between posterior samples. While TD mostly agrees with both rNBS and APD, there exists an anomalous region with high dissimilarity values based on TD at around iteration 6, 250. Upon inspection, we observe a minor topological change close to a pair of leaves, causing the topological dissimilarity value to peak at 7.0. While both rNBS and APD report a slight increase in dissimilarity in that region as well, the reported dissimilarity from neither measure is high enough to affect the dissimilarity scales of rNBS and APD. This highlights topological dissimilarity’s over-sensitivity to minor topological changes, which is not the case for either rNBS or APD.

### 3.3 Runtime Comparison

We report on the runtimes of the current implementations of rNBS, APD, and TD of [22] with respect to the number of reticulations in the pairs of networks. All experiments were run on a desktop running Linux Mint 20.3 on a single AMD Ryzen 9 5900X 12-Core Processor and 31.3 GiB available memory. Fig. 8 summarizes the results.

**Fig. 8:**
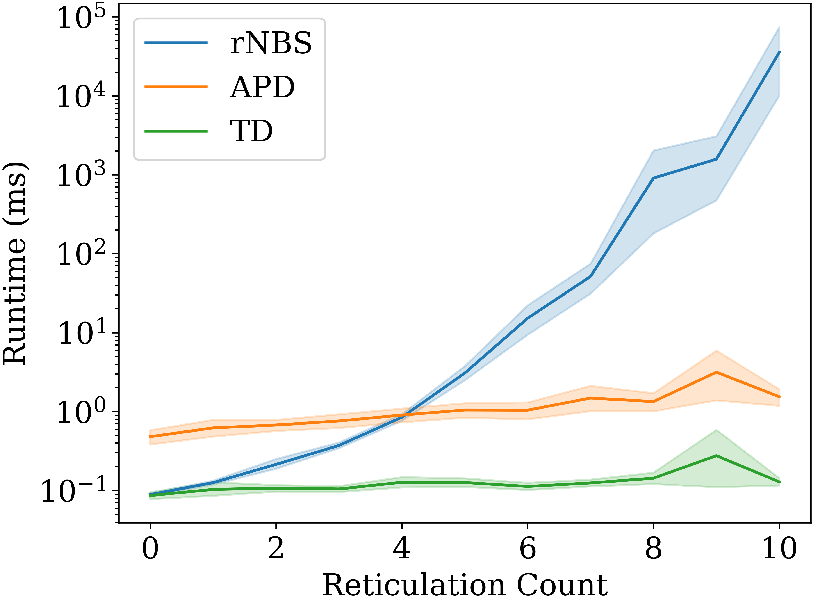
Runtimes of rNBS, APD, and TD of [22] with respect to number of reticulation nodes in the networks. Time is measured in milliseconds.

We find that APD is one fold slower than TD. Additionally, while the runtime increases as the number of reticulations increases, the rate of increase is very small. When there are no reticulations, APD can compute approximately 2,082 dissimilarities per second, and when there are 10 reticulations, APD can compute approximately 649 dissimilarities per second. In contrast, rNBS slows down significantly as the number of reticulations increases, which supports the asymptotic time complexity of the edge covering algorithms used for bipartite graphs formulated for rNBS (11, 278 per second when there are no reticulations vs. 0.03 per second when there are 10 reticulations). It should be noted that our implementation of APD is written in Java and does not currently use any libraries or hardware acceleration to transform matrices.

## 4 Conclusions and Future Work

By taking branch lengths into account, our two novel dissimilarity measures of phylogenetic networks will address an important deficit in the ability to analyse phylogenetic network space and its exploration. By taking different approaches to measuring dissimilarity, and through our analysis of scaled, perturbed, and MCMC sampled phylogenetic networks, we have shown that the path distance approach has more immediate promise than edge-covering of displayed trees. We implemented both measures in PhyloNet [31, 33] and studied their properties on pairs of perturbed networks. Furthermore, we illustrated the use of the dissimilarity measures for clustering and summarizing the phylogenetic networks in a posterior sample of the Bayesian inference method MCMC_SEQ. There are many directions for future work, two of which we will discuss here.

### Incorporating inheritance probabilities in the dissimilarity measures

Statistical inference of phylogenetic networks results not only in estimates of the topologies and branch lengths, but also of inheritance probabilities, which annotate the reticulation edges. The measures we presented above ignore inheritance probabilities. In particular, the APD measure treats all paths equally. One possible extension that accounts for inheritance probabilities is to weight each path that is counted by a combination of the inheritance probabilities used by the reticulation edges on that path. Thus, a further extension to average path distance can incorporate inheritance probabilities by weighting the path distance between two leaves *i* and *j* at each MRCA. Thus, for each MRCA, we multiply the path distance from the MRCA to leaf *i* with the inheritance probabilities along the path, perform the same for the path from the MRCA to leaf *j*, and sum them up. Finally, the weighted path distance (WPD) becomes the sum of weighted distances at each MRCA of *i* and *j*. For example, for the network of Fig. 4, the weighted path distance between *x*_2_ and *x*_3_ in the network is the sum of weighted path distances at the three MRCAs: [(λ_10_ + λ_6_ + λ_2_ + λ_1_) · *γ*_2_ + (λ_12_ + λ_8_ + λ_7_ + λ_4_)· (1 – *γ*_1_)] + [(λ_10_ + λ_6_ + λ_2_) · λ_2_ + (λ_12_ + λ_8_ + λ_5_) · λ_1_] + [(λ_10_ + λ_9_) · (1 – λ_2_) + λ_12_]. Similarly, for the rNBS measure, we can weight each tree by the product of the inheritance probabilities of the reticulation edges used to display the tree. However, it is important to note here that these measures could be very sensitive to inaccuracies of the inheritance probability estimates. For example, two networks that are identical in terms of topologies and branch lengths but vary significantly in the inheritance probabilities of even a small number of reticulation edges could result in a very large dissimilarity value, unless the inheritance probabilities are weighted carefully. We will explore these directions.

### Tree-based dissimilarity in the presence of incomplete lineage sorting

Zhu *et al*. [36] showed that when incomplete lineage sorting occurs, inference and analysis of phylogenetic networks are more adequately done with respect to the set of parental trees of the network, rather than the set of displayed trees. Thus, a different approach to defining the rooted network branch score could involve building a complete bipartite graph on the sets of parental trees of the two networks. A major challenge here could be computational. As we demonstrated in our analysis of rNBS, the runtime of the current implementation increases exponentially with the number of reticulations *k*. Worse, while a phylogenetic network has up to 2^*k*^ displayed trees, the number of parental trees could be significantly much larger to the extent that 2^*k*^ is feasible in the case of a small *k* whereas computing the set of all parental trees explicitly for the same value of *k* can be infeasible. Therefore, faster computations and/or heuristics for computing such a dissimilarity measure would be needed. As we discussed above, we will study whether constructing the bipartite graph explicitly is necessary for computing rNBS, as avoiding such construction could result in significant improvement to the computational requirements.

## Acknowledgements

We thank Zhen Cao for contributing the MCMC posterior sample files for the simulated data set. This work was supported in part by NSF grants CCF-1514177, CCF-1800723 and DBI-2030604 to L.N.

